# Rapid PCR-based screening system for detection of type II CRISPR-Cas loci in bacterial species

**DOI:** 10.64898/2026.08.24.746701

**Authors:** Ayesha Bibi, Tahir Iqbal, Kainat Ilyas, Asma Nosheen

**Affiliations:** Molecular Biology Lab, Department of Zoology, University of Gujrat, Pakistan; Institute of Zoology, University of the Punjab, Pakistan

**Keywords:** PCR, rapid screening, type II CRISPR-Cas, Cas9, bacteria

## Abstract

The Clustered Regularly Interspaced Short Palindromic Repeats (CRISPR) and associated nuclease gene (Cas), originating from the bacteria acquired immune system, have revolutionized gene editing technology. In this regard, type II (Cas9) been extensively studied and widely applied CRISPR system so far. The mechanism for precise manipulation of genomic sequences is guided by small RNA called CRISPR RNA (crRNA). In this study we devised and optimized CRISPR-Cas9 screening system based on Cas9 gene detection, targeting a conserved part of recognition domain (REC) consisting of arginine rich bridge helix (BH). We used hemi-nested PCR approach for screening sensitivity and reproducibility. The recombinant *E. coli* DH5 alpha containing the pRGEB32 vector (DH5 alpha/pRGEB32) with the Cas9 gene was used for system optimization. Subsequently, the screening system was applied and validated on different environmental bacterial strains including *Alcaligenes faecalis* and *Pseudomonas stutzeri*, isolated from sewerage samples. The optimized hemi-nested PCR resulted in amplification of targeted region in environmental bacterial strains and results were reproduced successfully. Furthermore, nucleotides and amino acid sequence, motif and domain analysis of PCR products, confirmed the targeted Cas9 REC-BH domain. Presently, no rapid and cost effective CRISPR-Cas screening system is available except expensive whole genome sequencing approach. Our investigation aimed to device rapid and cost effective screening system for identification of new variants of Cas9 proteins in environmental bacterial species. In this context, the developed Cas9 gene-based CRISPR-Cas screening system (C9CSS) may be a potential rapid screening tool to identify new Cas9 orthologs in different bacterial genomes with improved functions.

**IMPORTANCE:** The bacterial acquired immune system comprising CRISPR-Cas locus, has revolutionized targeted gene editing and now applications beyond it in the form clinical remedy including diagnostics and therapeutics for bacterial, viral and genetic diseases. Notably, Cas9 (Type II CRISPR) is the first one studied in detailed and programmed for different applications and discovery of its new orthologs with improved functions is still area of interest. But the limitation of large-scale screening of bacterial species for new orthologs identification due to expensive WGS workflow is a big challenge. To address this problem, we have developed and optimized a cost effective screening tool C9CSS to preliminary identify CIRISPR type-II locus in bacteria. Subsequently, only C9CSS positive sample may be processed for WGS for detailed characterization evading the cost of negative samples.

## INTRODUCTION

The Clustered Regularly Interspaced Short Palindromic Repeats (CRISPR) were discovered in *E. coli* and later on identified as a bacterial immune system against mobile genetic elements (MGEs), plasmids and phages (1,2). The CRISPR-Cas system keeps the bits of acquired genetic material as a memory which were encountered by previously pathogenic invaders. This system effectively defends against phage infections and plasmid conjugation (3). The defense mechanism facilitated by the CRISPR-Cas system is operated through three stages; first adaptation, the small DNA fragments incorporating as a new spacer into CRISPR array when a pathogen infects the bacterial cell (4), second expression, transcription occurs to produce a CRISPR RNA (crRNA) precursor located within a CRISPR locus which is further processed by breaking down the repeats to yield mature crRNA (5), third is interference, where a mature crRNA, a spacer surrounded by small DNA repeats, guides Cas protein to increase anti-viral response (6). The CRISPR-Cas system.is further divided into two classes; Class 1 with several effector proteins contains three subtypes (Type I, III, and IV) but Class 2 has single effector protein and three subtypes (Type II, V, and VI) (7). The CRISPR-Cas system can target DNA (Type I, Type II, and Type V), RNA (Type VI), and both DNA and RNA (Type III). As a general rule CRISPR Cas systems functional complex consists of effector protein with endonuclease activity, trans acting CRISPR RNA (tracrRNA) and crRNA which guides the targeted activity (8).

The Cas9 endonuclease belongs to type II system is extensively studied for it applications and is utilized for precise DNA targeting. In programmed version, the single guide RNA (sgRNA also shortly gRNA) formed by the combining the tracrRNA and crRNA as a single hybrid RNA molecule, which enabled the Cas9 to target different DNA target sites by changing the sgRNA sequences according to the target sites. The protospacer adjacent motif (PAM) identified by Cas9, plays a crucial role in facilitating the attachment and cleavage of DNA when guided by sgRNA-Cas9. A typical Cas9 protein consists of two lobes; one known as recognition (REC) and other a nuclease (NUC) lobe. The REC lobe is further subdivided into Bridge helix (BH) (aa 60–93) followed by REC1 (aa 94–179 and 308–713) and REC2 (aa 180–307) domain. Similarly, NUC lobe have RuvC (aa 1–59, 718–769 and 909–1098), HNH (aa 775–908), and PAM-interacting (PI) aa 1099–1368) (9). Both, REC and NUC lobes are connected by long arginine-rich BH domain and the disruptions of the BH-loop effect Cas9 activity at multiple levels. It is suggested that it interacts directly with sgRNA and facilitates the formation of binary complex of sgRNA and Cas9 protein further to execute targeted endonuclease activity (10). Similarly, the BH-RNA interaction was also reported in other types of CRISPR system namely Cas12a (11) and Cas12b (12).

The PCR based CRISPR-Cas rapid screening has been a big challenge and identification of bacterial strains containing native CRISPR-Cas system has been dependent on whole genome sequencing (WGS) and subsequent bioinformatics analysis to map CRISPR loci. In turn, the high-cost WGS limits large scale screening of bacterial isolates for the presence of native CRISPR-Cas system, as even negative samples must be sequenced in this scenario. In the present work we developed and optimized a cost-effective C9CSS PCR-based detection system for CRISPR-Cas9 system in bacterial strains. This screening system will be helpful as preliminary rapid detection tool to identify bacterial strains harboring CRISPR-Cas9 system which may be subsequently subjected to WGS and bioinformatics analysis for detailed classification and characterization. Hence, this two-steps screening approach reduces the overall cost and may expedite discovery of new Cas9 protein variants with improved performance.

## RESULTS

### Optimization

The optimization of C9CSS was performed using a hemi-nested PCR approach, with both rounds optimized independently. *E. coli* DH5 alpha/pRGEB32 was used as template bacterial strain for Cas9 gene amplification. In the first round 266 bp PCR product was obtained using outer primers set C9PF1 and C9PR1 (Fig. 1 A). While in second round 260 bp PCR product was amplified using a hemi-nested primers set C9PF1 and C9PR2 (Fig. 1 B). The optimized PCR amplification for both rounds was obtained on following conditions; initial denaturation at 95 °C for 5 minutes with 30 cycles of denaturation at 95 °C for 45 seconds, annealing at 50 °C for 30 seconds and extension at 72 °C for 50 seconds, followed by final extension at 72 °C for 7 min.

**FIG 1.**
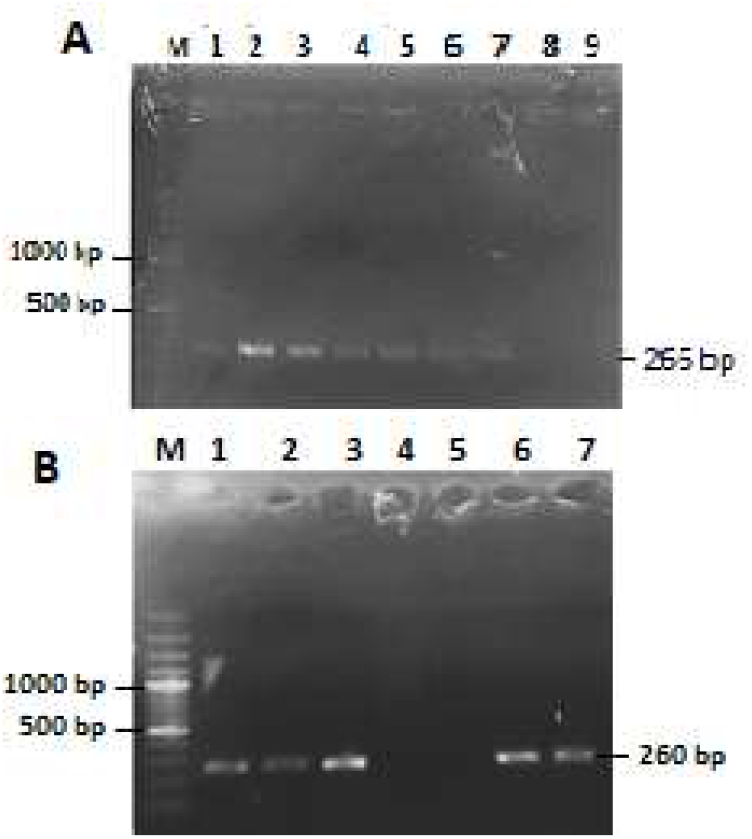
Optimization of C9CSS. **A)** first round hemi-nested first round PCR partial Cas9 gene amplification with primers (C9PF1, C9PR1) using *E. coli* DH5 alpha/pRGEB32 isolates templates. M= 100 bp DNA marker, Lane 1&2= M32A, Lane 3&4= M32B, Lane 5&6= M32C, Lane 7& 8 = S1, lane 9 = negative control. **B)** second round hemi-nested PCR amplification of partial Cas9 gene (primer set: C9PF1, C9PR2) using *E. coli* DH5 alpha/pRGEB32 isolates template. M= 100 bp DNA marker, Lane 1= M32A, Lane 2= M32B, Lane 3= M32C, Lane 4 and 5= negative control, Lane 6= S1, Lane 7= positive control.

### Application to environmental bacterial strains and reproducibility

Followed by optimization of the C9CSS, it was applied to previously identified bacteria from sewerage samples *Alcaligenes faecalis* (GTA and GTC) and *Pseudomonas stutzeri* (1ACG, 2ACG, 3ACG) and *E. coli* DH5 alpha/pRGEB32 was used as positive control. All samples showed successful PCR amplification in both first (Fig. 2 A) and second (Fig. 2 B) round with required product size. Subsequently, reproducibility of the C9CSS was analyzed through a typical hemi-nested PCR approach using PCR product of the first round as template for second round amplification with each sample processed in duplicate. The 260 pb hemi-nested PCR product was obtained in all reactions (Fig. 3).

**FIG 2.**
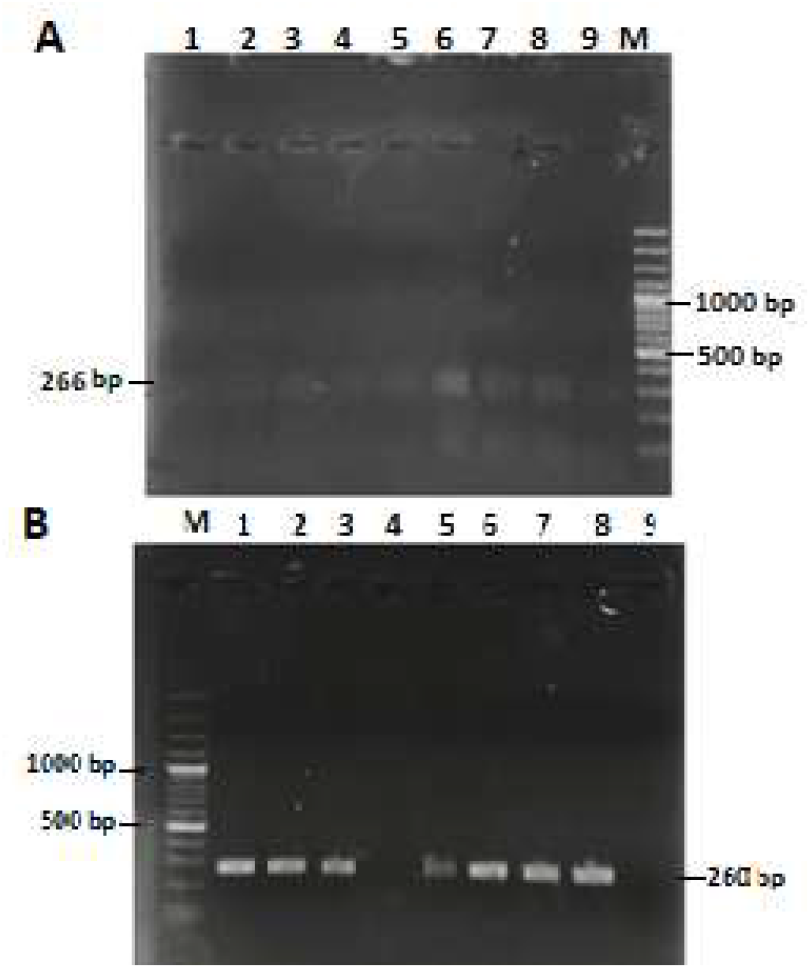
Application of C9CSS to bacterial strains isolated from sewerage samples. **A)** first round hemi-nested PCR using primers C9PF1, C9PR1. M = 100 bp DNA marker, Lane 1&2 = negative control, Lane 3 = *E. coli* DH5 alpha/pRGEB32 (positive control), Lane 4 = S1 (positive control), Lane 5 = GTA, Lane 6 = GTC, Lane 7 = 1ACG, Lane 8 = 2 ACG, Lane 9 = 3 ACG. **B)** second round hemi-nested PCR with primers C9PF1, C9PR2. M = 100 bp DNA marker, Lane 1 = *E. coli* DH5 alpha/pRGEB32 (positive control), Lane 2 = S1 (positive control), Lane 3 = GTA, Lane 4 = negative control, Lane 5 = GTC, Lane 6 = 1ACG, Lane 7 = 2 ACG, Lane 8 = 3 ACG, Lane 9 = negative control.

**FIG 3.**
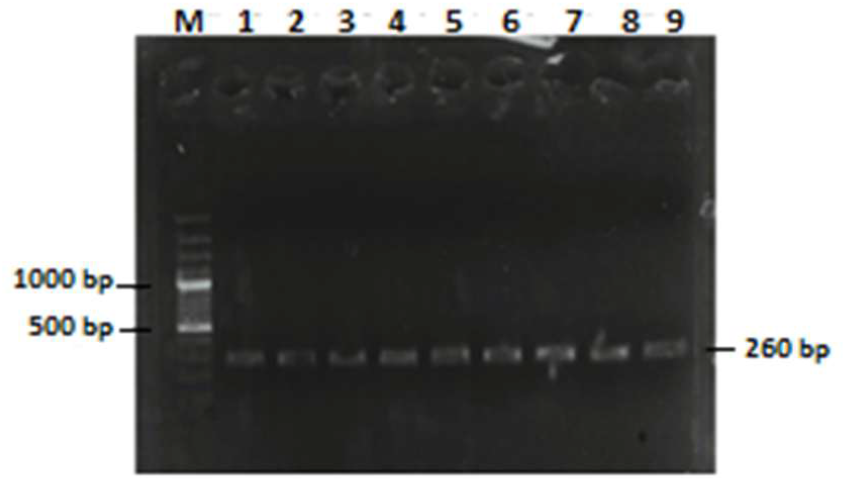
Reproducibility of C9CSS – two steps hemi-nested PCR amplification of partial Cas9 gene. Lane M = 100 bp DNA marker, Lane 1&2 = GTA, Lane 3&4 = GTC, Lane 5&6 = 1ACG, Lane 7&8 = 2ACG, Lane 9 = 3ACG.

### Sequence and domain analysis

After BLAST analysis the sequences of the PCR products were subjected to multiple sequence alignment which showed conserved regions highlighted in red boxes in figure 4. The motif and their location analysis revealed presence of seven motifs ranging from 14 to 50 nucleotides (Fig. 5 A). Motif 1, 2, 4 and 5 were found conserved in all bacterial strains. Notably, among these conserved motifs, motif 1 showed the most conserved sequence with 98 % identity followed by motif 2 with 64 % sequence identity among studies bacterial isolates (Fig. 5 B). The obtained amino acid sequence, nucleotide sequence of each sample was translated (Figure 6 A) and the amino acids multiple sequence alignment showed conserved regions, particularly arginine rich BH in the REC domain (Fig. 6 B). This was further reconfirmed by protein domain and Pfam analysis which showed a significant hit to the Cas9-BH in REC domain (PF16593) with highly reliable E-value 1.6 × 10−^10^, spanning amino acid position 29 – 57 (Fig. 7 A and B). The BH has critical role in guide RNA interaction and nuclease activity of Cas9. The second match to mRNA triPase domain (PF02940) – mRNA capping enzyme, was detected with relatively high E-value 0.029 and amino acid positions 8-57. Due to low confidence score and functional irrelevancy with Cas9, this domain was not considered. Notably, the phylogenetic analysis of the strains on the basis of Cas9 BH domain showed the strain GTC and 1ACG most divergent among the studied strains (Fig. 8).

**FIG 4.**
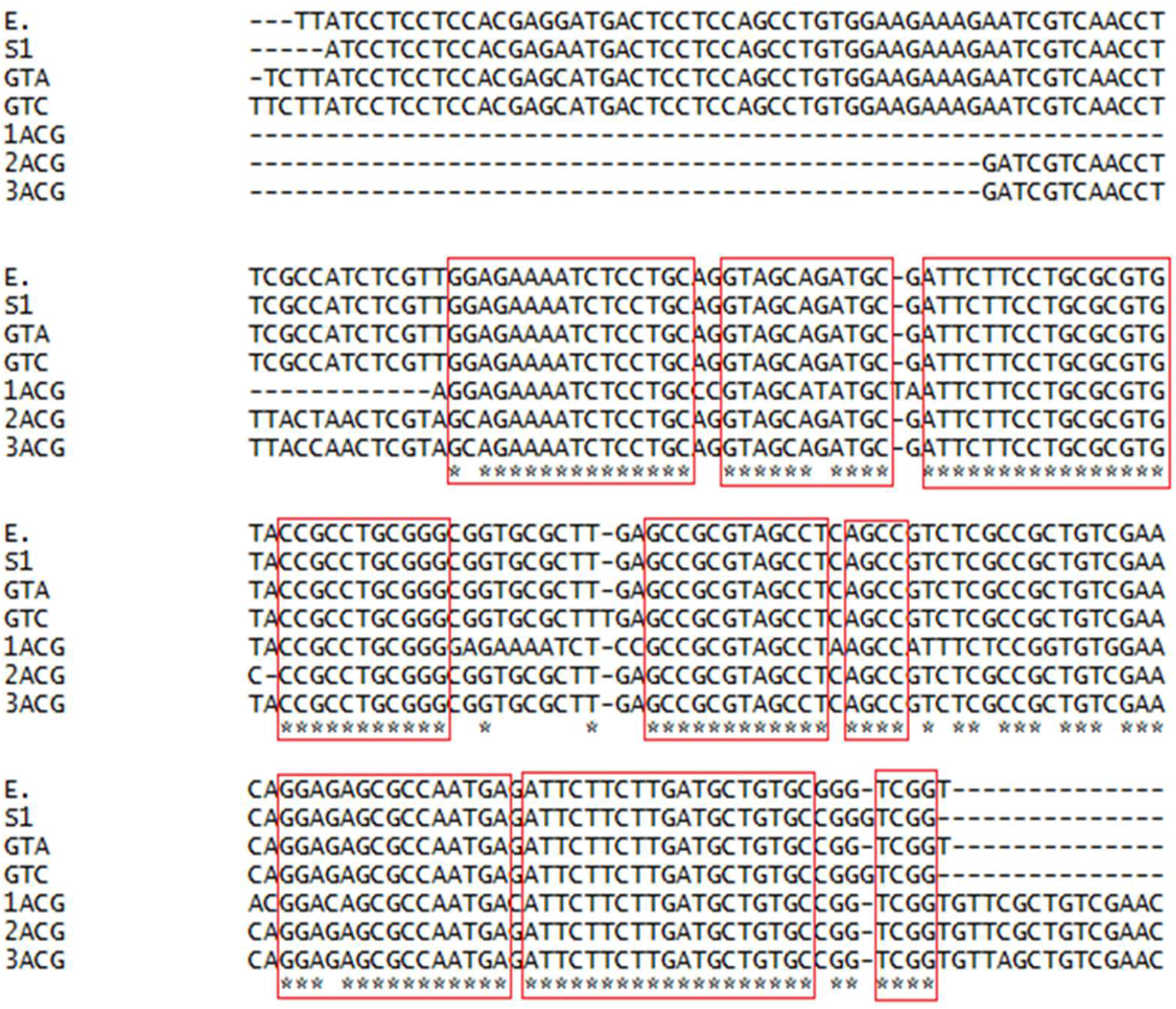
Multiple sequence alignment (MSA) of partially amplified Cas9 gene REC domain (BH)

**FIG 5.**
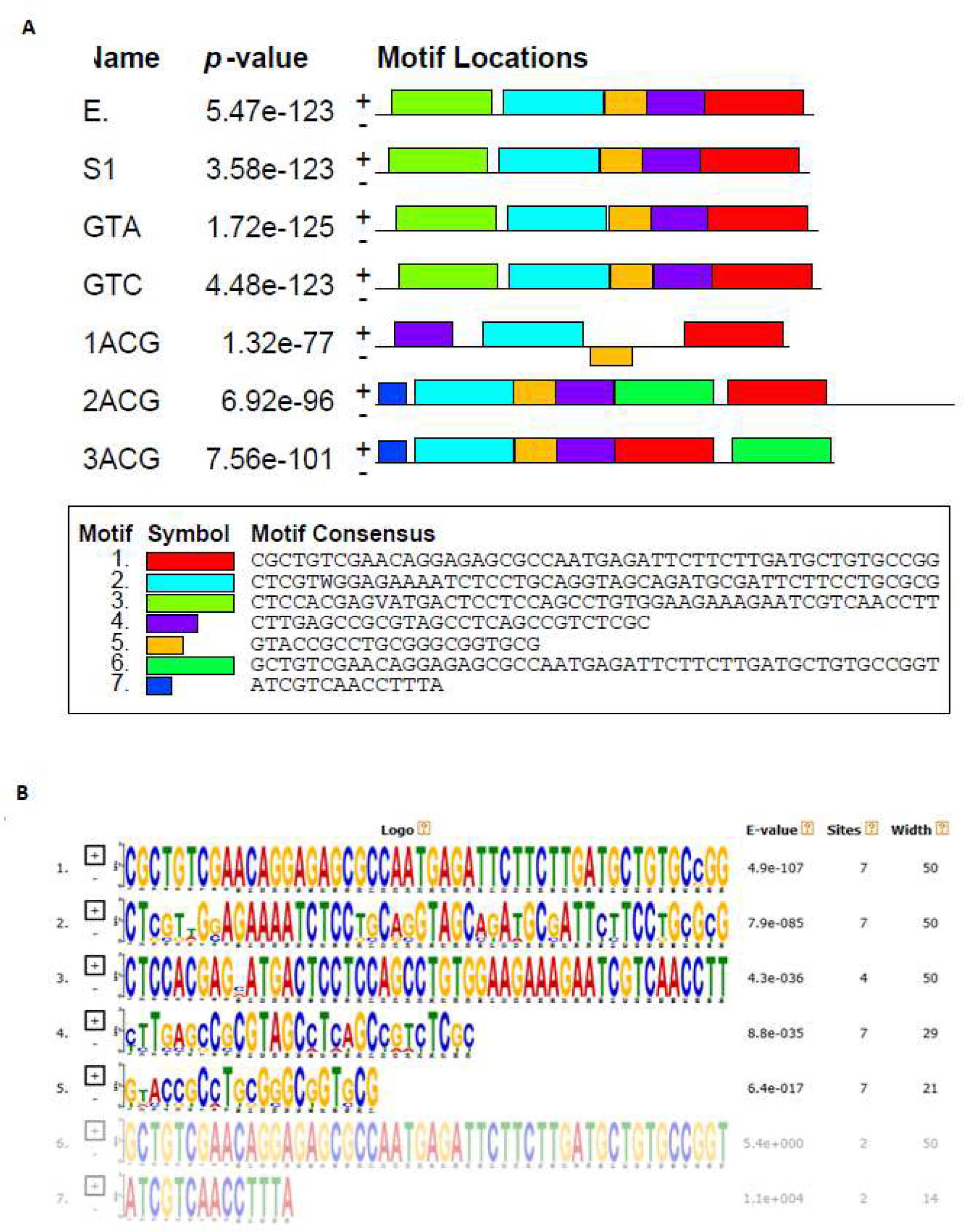
Nucleotides sequence motif analysis. **A)** Motif analysis of the Cas9 gene amplified region.**B)** Sequence homology of identified motifs among the studied bacterial strains.

**FIG 6.**
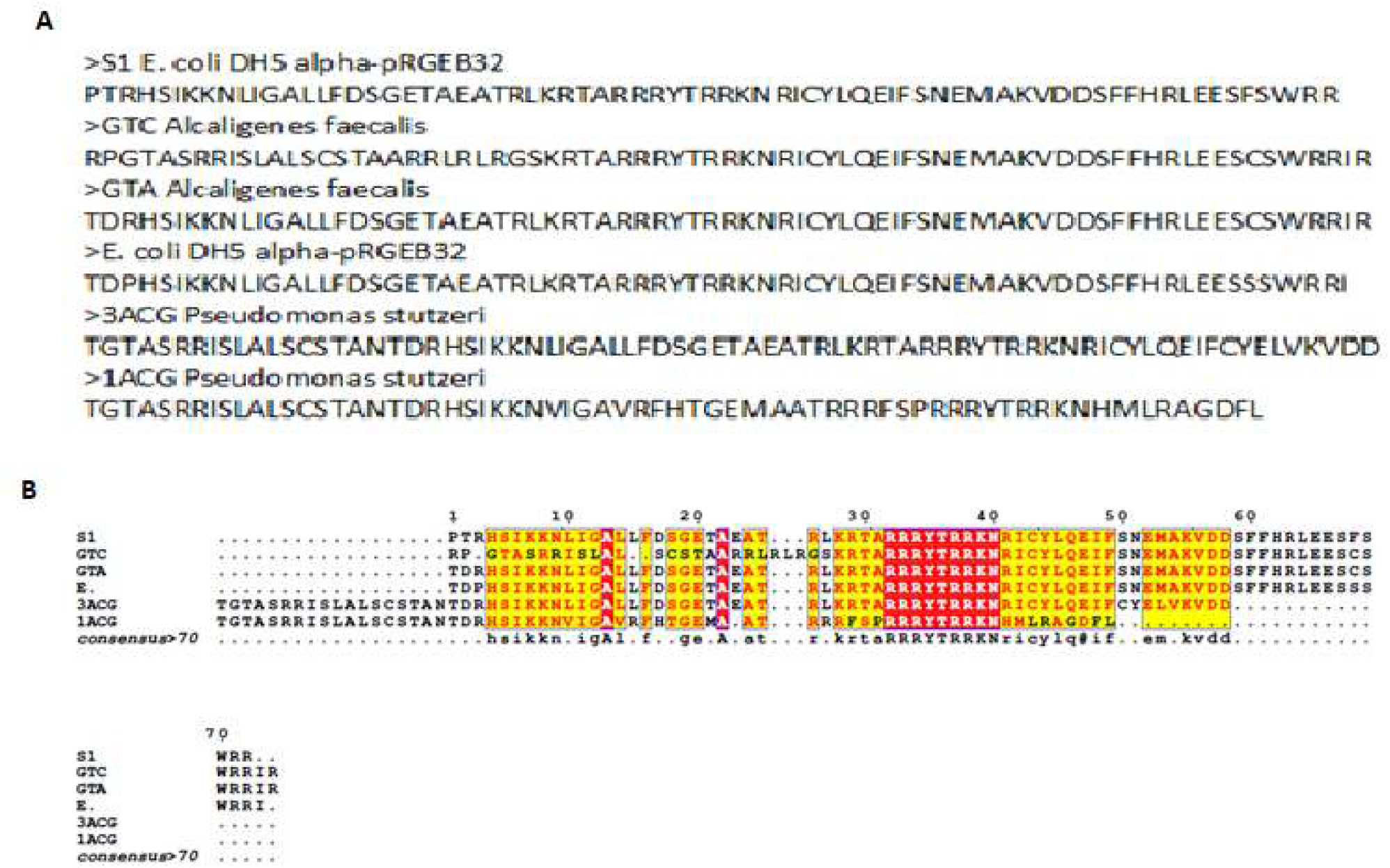
Amino acid sequences of amplified Cas9 gene portion of bacterial strains. **A)** Translated amino acid sequence. **B)** Amino acid sequence based multiple sequence analysis of amplified partial Cas9 RCE domain (BH region).

**FIG 7.**
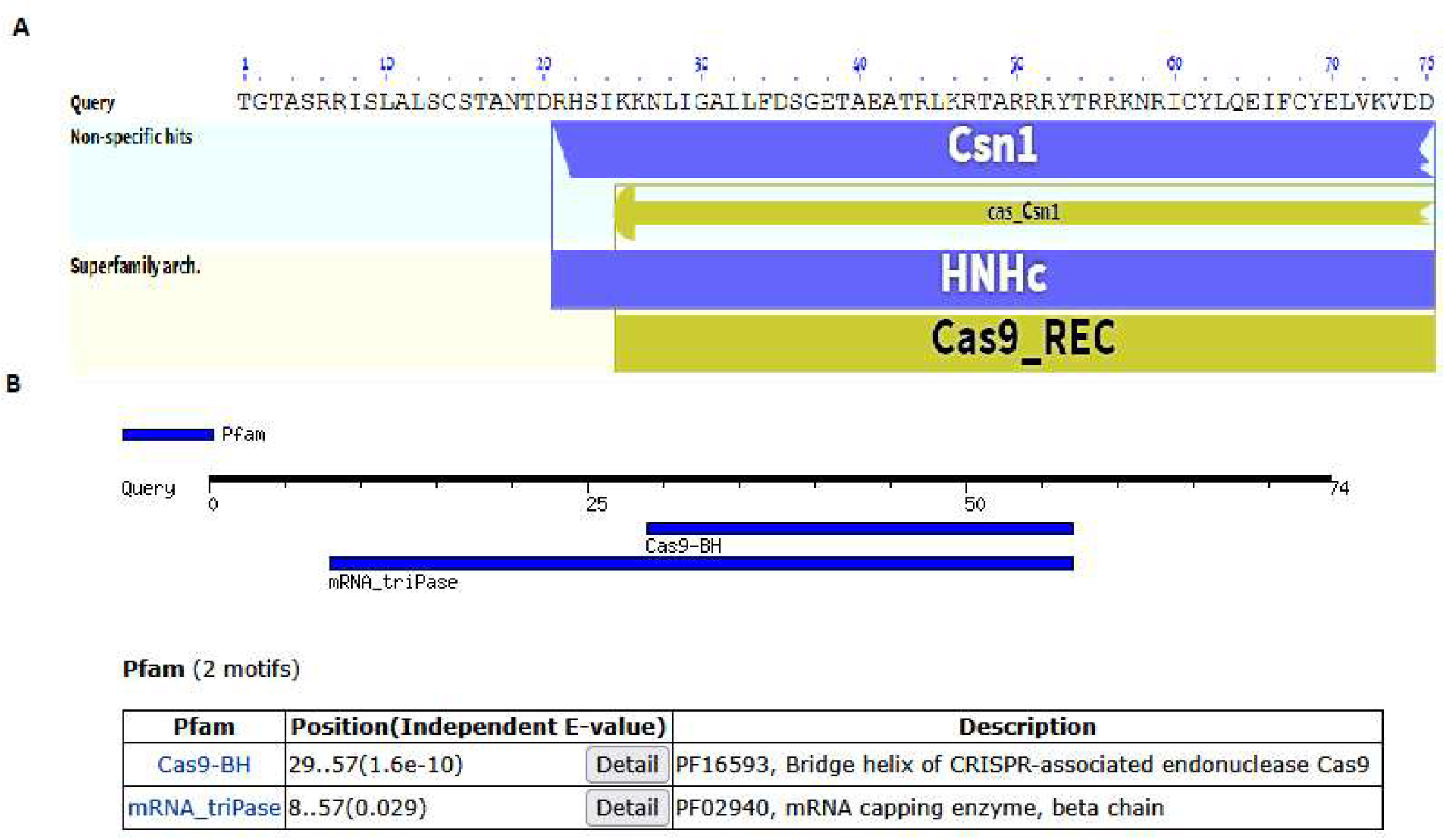
Protein identification analysis of PCR amplified region. **A)** Protein domain analysis. **B)** Protein family motif analysis.

**FIG 8.**
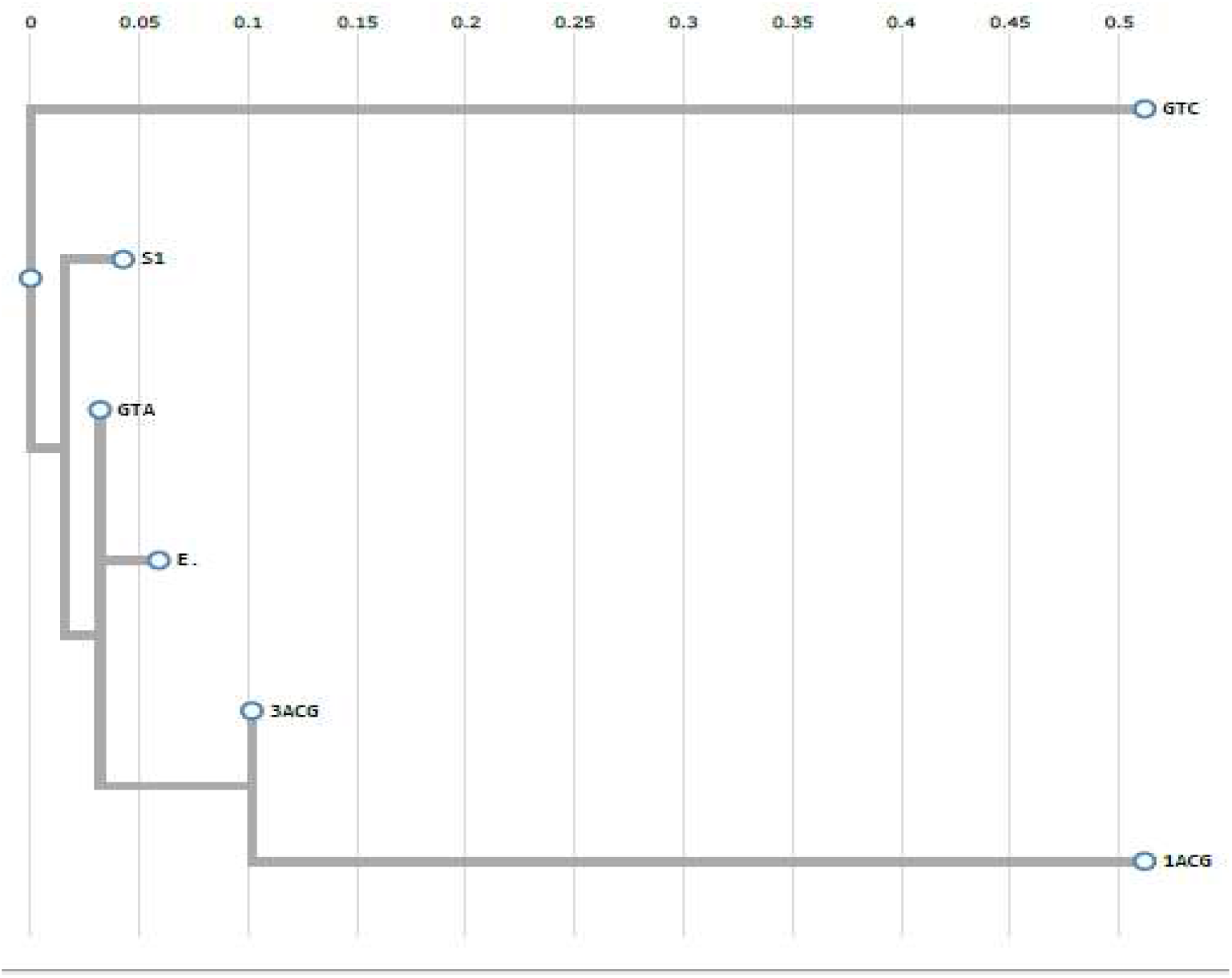
Phylogenetic relationship of bacterial strains on the basis of Cas9 partial nucleotides sequence of REC domain BH region.

## DISCUSSION

Rapid screening of CRISPR-Cas system in bacterial species has been a big challenge. So far the only way seems to be the whole genome sequencing and afterward analysis of complete genome sequence data using *in silico* tools to detect CRISPR-Cas locus. The genetic variations pose main limitation for developing a universal rapid screening system based on PCR technique. We designed and developed a rapid C9CSS hemi-nested PCR based screening system targeting of Cas9 gene portion encoding conserved BH –domain. Hence, C9CSS may be used as a universal PCR based screening system for detection of Type-II CRISPR locus in different bacterial strains. The wide range applications of Cas9 in biological fields drive the screening of more efficient orthologs for targeted clinical utilization (13).

The of BH-domain is critical for Cas9 cleavage activity and mutations result in impaired nicking of DNA molecule and lack of target linearization in case of circular DNA molecules (10). Keeping in view the primary role, structural and functional conservation of BH-domain of Cas9 protein, we devised C9CSS a universal PCR based detection, to minimize the effect of genetic diversity, of type-II CRISPR locus among bacterial strains. subsequently, it was evident from our study by demonstrating the detection of Cas9 gene in *Alcaligenes facecalis* and *Pseudomonas stutzeri* isolated from sewerage samples. The sewerage environment has abundant and diverse bacterial communities and accordingly wide range of phages thrive this environment (14, 15). Due to frequent interaction and excessive phages prevalence may drive the development of active CRISPR-Cas9 locus as combat immunity in these bacteria. In this context, we applied C9CSS to sewerage isolated bacteria to assess validation and system functionality.

The target verification by nucleotide and amino acid sequence analysis of PCR products showed detection of Cas9 BH-domain and nucleotides motifs. Out of total 7 predicted nucleotides motifs, 1 and 2 were found conserved among all studied bacterial strains suggesting signature sequence markers for Cas9 gene (Figure 7). Furthermore, domain and protein family motif analyses of amino acid sequences exhibited Cas9 REC (BH) domain recognition which is the conserved functional part of Cas9 protein and may be helpful in detection of different Cas9 orthologs in different bacterial strains (16, 11). Worth noting, the detection of signature BH-domain with arginine rich residues sequence in each sample showed sensitivity and reproducibility of the results (10). Furthermore, the hemi-nested PCR ensured sensitivity by two round targeting; in first round a large fragment is amplified and the second round amplified smaller fragment nested within it. The use of hemi-nested PCR, due to its specificity and sensitivity, for detection of bacterial and viral genome or specific gene (s) as diagnostic tool is well established (17).

In summary, the hemi-nested PCR based C9CSS screening system was devised for universal detection of type-II CRISPR locus by targeting Cas9 gene. It specifically targets BH-domain which is the conserved part of Cas9 gene. The C9CSS was successfully applied to *Alcaligenes facecalis* and *Pseudomonas stutzeri* isolated from sewerage waste.

## MATERIALS AND METHODS

### Bacterial strains culturing and identification

The recombinant *E. coli* DH5 alpha/pRGEB32 was used in this study, kindly gifted by Center of Excellence in Molecular Biology (CEMB) University of the Punjab. The pRGEB32 vector (Addgene) contains Cas9 gene with kanamycin resistance gene as selectable marker for recombinants selection and we use it as a template for amplification of Cas9 gene to optimize C9CSS rapid screening system. The *E. coli* DH5 alpha/pRGEB32 strain was cultured on kanamycin (50 μg/mL) LB selective medium.

The previously isolated and molecular identified sewerage bacterial strains were used in this study for validation of C9CSS. These bacteria were isolated from sewerage samples collected from main sewerage lines of Gujrat city, Punjab, Pakistan (32.573770, 74.076897) (unpublished data).

### Designing and development of Cas9 gene based CRISPR detection system

The C9CSS tool was designed using hemi-nested PCR approach to detect Cas9 gene in bacterial strains. Accordingly, the primers were selected from conserved REC domain of Cas9 gene, identified by multiple sequence alignment (MSA) using ClustalW (18). The developed hemi-nested PCR detection system consisted of three primers; outer primer pair (first round), C9PF1 forward primer 5’-AACACCGACCGCACAGCATCAAGAA-3’, external reveres primer C9PR1 5’-TCGTCCACGATGTTGCCGAAGA-3’, and internal reverse primer, C9PR2 5’-ACGATGTTGCCGAAGATGGGGT -3’, pairing with C9PF1 (first round forward primer) for final round PCR. The final PCR product size was 260 bp.

### Optimization of Cas9 screening system

The optimization of C9CSS tool was carried out using recombinant *E. coli* DH5 alpha/pRGEB32 as template bacterial strain. Two round hemin-nested PCR was done using primer set C9PF1/ C9PR1 in first round and C9PF1/ C9PR2 in second round. The annealing temperature ranging 45-55 °C was screened to obtain optimum annealing temperature. So, overall PCR conditions used for both rounds were as under; initial denaturation 95 °C for 5 minutes followed by 30 cycles of denaturation at 95 °C for 45 seconds, annealing range 45 - 50 °C for 30 seconds and extension at 72 °C for 50 seconds. Final extension was completed at 72 °C for 5 minutes. Furthermore, template negative control was used to ensure no template contamination in PCR reagent. Subsequently, obtained PCR products were subjected to horizontal gel electrophoresis with 1.5% agarose gel.

### DNA sequencing and sequence analysis

The PCR products were sequenced through Sanger sequencing method. Subsequently, the obtained sequences were subjected to Basic Local Alignment Search Tool (BLAST) analysis for verification and retrieval of similar sequences to proceed further for phylogenetic analysis. In the next step the multiple sequence alignment was done using CLUSTAL W tool (18) and phylogenetic analysis was done using PhyML tool (19). Motif mapping was done through MEME suit 5.5.9 (20). Further, the nucleotide sequences were translated using Expasy translate tool and the protein sequences were subsequently analyzed through protein-protein BLAST (blastp) for identification.

## ACKNOWLEDGEMENTS

We acknowledge the University of Gujrat for basic funding for this research work. We further acknowledge Center of Excellence in Molecular Biology (CEMB) University of the Punjab for generous gift of pRGEB32 vector (Addgene). This research received no specific grant from any funding agency in the public, commercial, or not-for-profit sectors.

